# Using landscape biodiversity metrics to assess rewilding: a space-for-time comparison between the Knepp Estate and an agricultural baseline at Boothby Wildland

**DOI:** 10.1101/2025.01.16.633386

**Authors:** NJ Pates, BB Boyce, MJ Crawley, Ivan de Klee, Penny Green, Josh A Hodge, Catalina Estrada, Eimear K Loughlin, Rona Learmonth, Thomas J Moye, Saskia IL Pearce, Megan Sherlock, William D Pearse

## Abstract

1. The biodiversity crisis is often framed in terms of the reduction of species found at a site: a (alpha) diversity. However, changes in species composition across a landscape – b (beta) diversity – caused by biotic homogenisation are part of the same crisis. Much biotic homogenisation in British terrestrial landscapes in the post-war period (1945 onwards) has been driven by agricultural conversion and intensification. Restoration efforts must therefore contend with restoring agricultural land to some desired state, and rewilding has emerged as a potential solution to this problem.
2. Here we quantify rewilding success by comparing a- and b-diversity of plant assemblages across two study systems in different stages of rewilding. Boothby Wildland is an arable farm recently given over to rewilding, while the Knepp Estate is an ex-arable and dairy farm currently >20 years into rewilding. We assessed a-diversity within, and b- diversity across, these two landscapes using 3 years of plant survey data (2022-2024).
3. We confirmed expected differences between a baseline and rewilded landscape and explored the changes in a brand new rewilded landscape in its early years of progress. As expected, the plant community at the Knepp Estate is relatively stable while that of Boothby Wildland is changing rapidly, and we discuss the landscape impacts of different herbivory regimes on recovery.
4. *Synthesis and applications.* We also show that spatial metrics for SESMPD and SESMNTD can distinguish between a rewilded and degraded landscape. This suggests that changes in the structure of biodiversity across landscapes can be quantified using analyses that account for spatial structure in metrics. We call for further research to see if a spatial approach of this kind can be used as a general metric of success in rewilding.

## Introduction

The biodiversity crisis is one of the most serious threats facing the planet (Pörtner et al., 2021). It is often defined as a reduction of α-diversity – the loss of local, site-level diversity (Newbold et al., 2015) – but loss of β-diversity – variation among sites – is at least equally significant (Newbold et al., 2018, Gossner et al., 2016). Loss of β-diversity, or biotic homogenisation, is the reduction of genetic, taxonomic, and functional diversity across landscapes (Olden et al., 2004, Olden and Rooney, 2006, Clavel et al., 2010, Gámez-Virués et al., 2015). Agricultural intensification is a major driver of biotic homogenisation in terrestrial systems (Sirami et al., 2019, Gossner et al., 2016) as large areas are merged into crop monocultures, native habitats are cleared, and water regimes are changed. This leads to decreased genetic and spatial diversity of local species (McKinney and Lockwood, 1999, Olden et al., 2004) together with reduced total taxonomic diversity. Agricultural expansion and intensification have been major themes of land-use change across Europe since the end of the second world war (Robinson and Sutherland, 2002) and an estimated 1.9 million km^2^ of land worldwide has been converted to agriculture since 1960 (Winkler et al., 2021). The pace of land conversion is still rapid, and demand for agricultural productivity is set to increase over the coming decades (Zabel et al., 2019).

The United Kingdom (UK) is an extreme example of habitat loss (Burns et al., 2023). A relatively small landmass with large population centres, 72% of UK land is agricultural (Hayhow et al., 2019). Since the chance to conserve most wild spaces in the UK has been missed, landscapes must now be restored and then protected. Ecological restoration through rewilding may offer solutions to the biodiversity crisis (Bailey et al., 2022, Environment Agency Chief Scientist’s Group, 2022). Rewilding has been part of the ecological literature for roughly 30 years and has been used in many contexts during that period (Johns, 2019, Schulte to Bühne et al., 2022). However, there is no fixed definition (Corlett, 2016a, Gann et al., 2019) and a lack of global agreement persists about the best way to conduct rewilding efforts (Suding et al., 2015). Rewilding has gained mainstream recognition in recent years (Jørgensen, 2015, Pettorelli et al., 2019), and is an increasingly popular option for projects seeking to recover and protect biodiversity (Donadio, 2022). This rise in popularity may be linked to the many proposed benefits of rewilding, including economic (Corlett, 2016b), human health and wellbeing (Maller et al., 2019), and landscape resilience (Lorimer et al., 2015) considerations. Despite the potential benefits, implementing rewilding is not simple. ‘Wild’ means different things to different groups (Ward, 2019), and so defining success in rewilding is challenging – a problem which is shared with traditional ecological restoration approaches. Targets for rewilding may use deep historical benchmarks (Donlan, 2005, Lorimer et al., 2015), invoke the concept of novel ecosystems (Corlett, 2016a, Hobbs et al., 2009), or use local remnants of undisturbed habitat as reference states (Corlett, 2016a, Gann et al., 2019). Others conceptualise rewilding not as a means to a defined end-state (Durant et al., 2019) but as a way to set landscapes on a trajectory that restores ecosystem function while requiring limited human intervention without a target ecosystem state in mind (Schweiger et al., 2019, Torres et al., 2018, Pettorelli et al., 2019). The latter definition is used here.

Because of extensive habitat loss and nature depletion, another problem that rewilding must contend with is that species that have historically contributed to ecosystem function may be missing from the landscape or present only in small populations. Globally, terrestrial systems have been radically impacted by land use change in recent decades (Winkler et al., 2021), but humans have been impacting systems for millennia. Many species have been lost since the late Quaternary (Sandom et al., 2014, Ripple et al., 2019), particularly large herbivores (Ripple et al., 2015, Smith et al., 2018). Loss of these species has consequences for ecosystem function: loss of connectivity and dispersal; changes in vegetation and soil structures; disruption of nutrient, gene, and energy flows; and changes in fire regimes (Berti et al., 2020, Cromsigt et al., 2018). Reintroduction or replacement of lost species, especially herbivores, is therefore often a key component of rewilding (Schweiger et al., 2019, Cromsigt et al., 2018, Svenning et al., 2016). However, if the goal is to achieve a trajectory and restore ecosystem services, we should not place too much focus on target species. Instead, we need metrics that will help us to understand when a system is moving in the right direction. Metrics need to be relatively simple to survey for if they are to be scalable and cost-effective, and they need to tell us something about the structure of the system as a whole – ideally from macro to micro scale and incorporating multiple biotic and abiotic systems as appropriate. Lack of spatial consideration is common in the evaluation and monitoring of restoration projects (Planchuelo et al., 2020, Tambosi et al., 2014) and, we argue, reduces how effectively a landscape’s recovery can be assessed. We propose that the required solution is a methodology for monitoring rewilding projects over time that focuses not only on the local diversity of a site (α-diversity) but also on how that diversity is arranged across the landscape (β-diversity). If a homogenous agricultural baseline is well-characterised, sites can then be assessed as having progressed from a state of biotic homogenisation by examination of the structure of their diversity in comparison with the baseline.

Here we used spatial analyses of diversity metrics to assess the structure of two test landscapes. We took a suite of α-diversity metrics and analysed the partitioning of variance within each metric at multiple spatial scales across our case study sites. In this way we studied variation in community structure (Anderson et al., 2011) to capture how landscape configuration affects diversity. We tested our method in a case study comparing 3 years of data collected at the Knepp Estate – the oldest and best-known example of rewilding in the UK – with Boothby Wildland – a baseline agricultural landscape that began rewilding in 2023. Both systems were recovering from the effects of intensive agricultural practices, but Knepp was more than 20 years into restoration while Boothby just beginning an exit from farming. Our goal was to find metrics that were able to (1) detect differences in structure between rewilded and agricultural systems (Knepp vs. Boothby in 2022) and (2) identify when the trajectory of rewilding is on course (Knepp vs. Boothby 2022-onwards). We demonstrate that this method can be used to show and quantify progress in rewilding projects.

## Materials and methods

We collected plant survey data at two locations in England to compare how biodiversity metrics are affected by changing heterogeneity of landscape across multiple spatial scales. We used a nested fractal design to select the 1m^2^ sites to survey at each location (Simpson and Pearse, 2021) (Figure 1A-C) and conducted plant surveys over a 4-week period in June-July in each survey year: 2022, 2023, and 2024.

**Figure 1.**
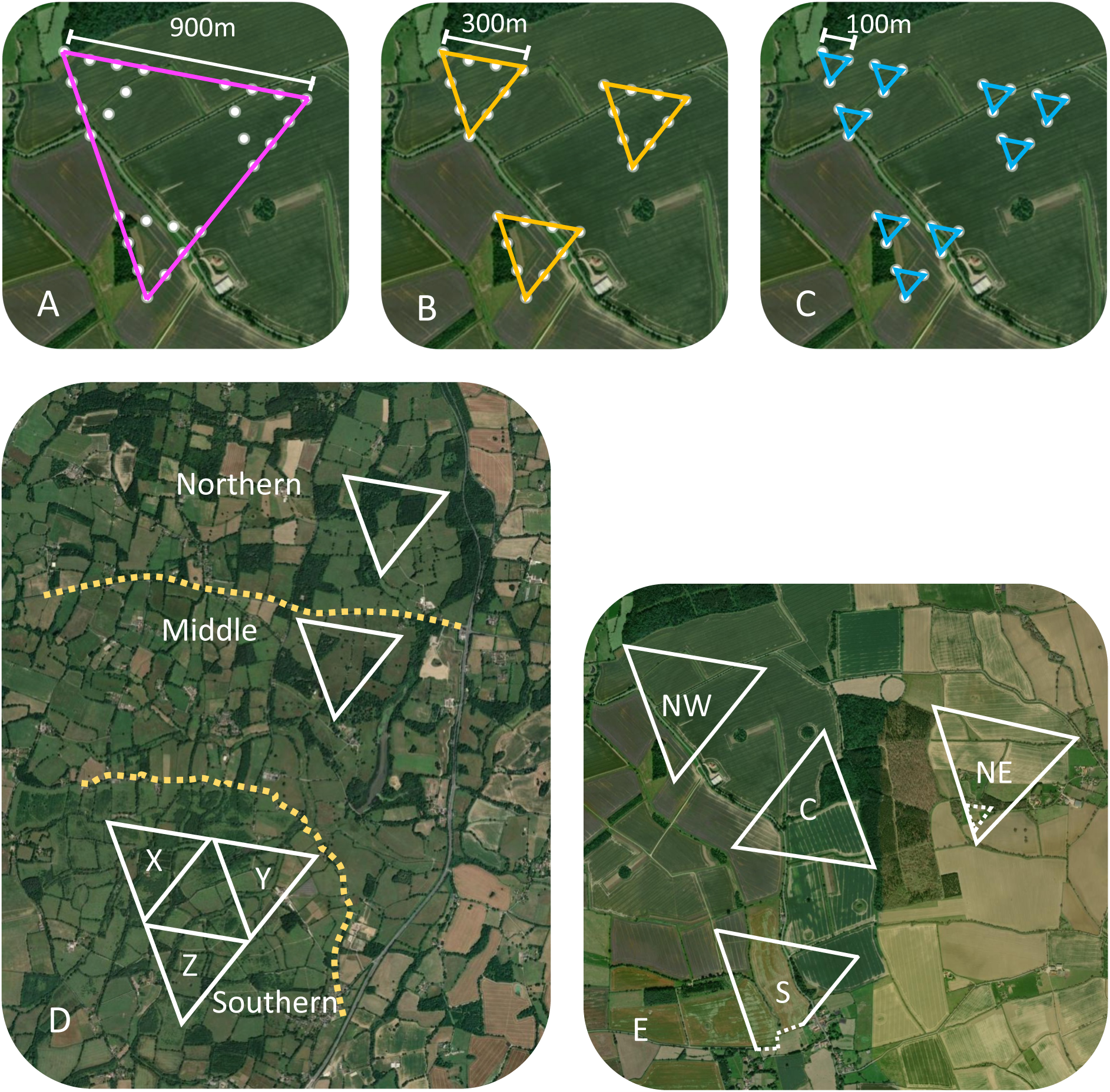
Nested fractal survey design: a flexible methodology to study landscape-level variation in biodiversity. **A.** A **fractal** is composed of 27 sites, and fractal-level grouping incorporates data from all 27 (pink triangle). **B.** Each fractal has three **major triads** of nine sites each (yellow triangles). **C.** Each major triad is comprised of three **minor triads** of three sites each (blue triangles). This survey design can be placed over any terrestrial landscape, arranged to maximise surveyable area i.e. avoiding large water features / out of bounds areas. Multiple fractals may be placed to maximise the area surveyed and account for irregular shaping in landscape boundaries. Here, the basic design (A-C) was adapted for the three differently sized parcels of land at Knepp (D) and land boundaries at Boothby (E). **D.** Survey design for Knepp. Knepp is divided into three blocks: northern, middle, and southern (separated by roads = yellow dotted lines). One fractal was placed in the northern block and one in the middle block. The southern block was large enough to accommodate three intersecting fractals to maximise the area surveyed: X, Y and Z. **E.** Survey design for Boothby. 4 fractals were positioned (NW, C, NE, and S). Dashed white lines indicate property boundaries.

### System histories

We surveyed two systems: the Knepp Estate (Knepp) and Boothby Wildland (Boothby). Knepp is a 3,500-acre ex-arable farm in West Sussex (50.97194, −0.36373) that has been rewilded since 2001 (Kirby, 2020, Tree, 2018). Knepp is split into three blocks (northern, middle, and southern) (Figure 1D). Each block has been subject to different grazing regimes following introductions of large herbivores (Kirby, 2020). Boothby, in Lincolnshire (52.86135, −0.54432), was still a 1,500-acre active arable farm in 2022 (Figure 1E). It comprised a patchwork of crop fields separated by hedgerows and patches of unmown field edges with small pockets of woodland present. Rewilding began in 2022-23, with approximately one third of the land being turned over to rewilding in each 2022-23 and 2023-24. These two systems are ideal to investigate patterns of biodiversity with changing landscape heterogeneity across multiple spatial scales: Knepp is one of the oldest lowland arable examples of a rewilded system in the UK whereas Boothby provides a valuable comparative agricultural baseline at “Year 0”. Year 1 and Year 2 data from Boothby then provide an indication of how quickly changes in community structure may be expected to occur in rewilding landscapes of this type. Agricultural land selected for rewilding in the UK will likely resemble Boothby structurally, and while the context of rewilding projects will vary, we argue that progress at Knepp and Boothby can be used as a guide for how increasing landscape heterogeneity can be linked to changes in diversity.

### Data collection

Following Simpson and Pearse (2021), we placed a 1m^2^ quadrat at each site surveyed. We recorded the species identity and percent coverage of vascular plant species rooted within the plot. Cover was estimated by eye to the nearest 1% for each species, unless cover was <1%, in which case an estimate to the nearest 0.1% was used. Species were identified in the field using guides or later from samples and photographs with additional resources (Cope and Gray, 2009, Rose, 2006). Because the habitat at Boothby comprised predominantly monocultures of cash crops in 2022 and 2023 (wheat, barley, broad beans, and oil-seed rape) we used an extended methodology there to allow us to include the maximum possible number of sites. We surveyed several sites in each crop field following the full protocol, then assumed that remaining sites in the same fields were similarly populated and confirmed a sub-set of these by photographing the location (see supporting information). To ensure that this extended methodology did not inflate the degree of homogenisation we recorded, we validated our approach by repeating our analysis using the full data set, and with the assumed and photographed sites removed. The results of our analyses were qualitatively identical, supporting this approach (Figure A1.1 and Figure A1.2). In 2024, when the landscape was considerably less homogenous, all sites were surveyed using the full methodology.

### Biodiversity metrics

We used four α-diversity metrics to investigate the effects of rewilding following Simpson and Pearse (2021): species richness (SR), Faith’s phylogenetic diversity (Faith, 1992) (Faith’s PD), standardised effect size of mean pairwise distance (SES_MPD_), and standardised effect size of mean nearest taxon distance (SES_MNTD_) (Figure 2). This spread of metrics allowed us to inspect both taxonomic and phylogenetic aspects of biodiversity (Tucker et al., 2017). The full data set and code to reproduce statistical analyses are provided online (DOI: 10.5281/zenodo.14035907).

**Figure 2.**
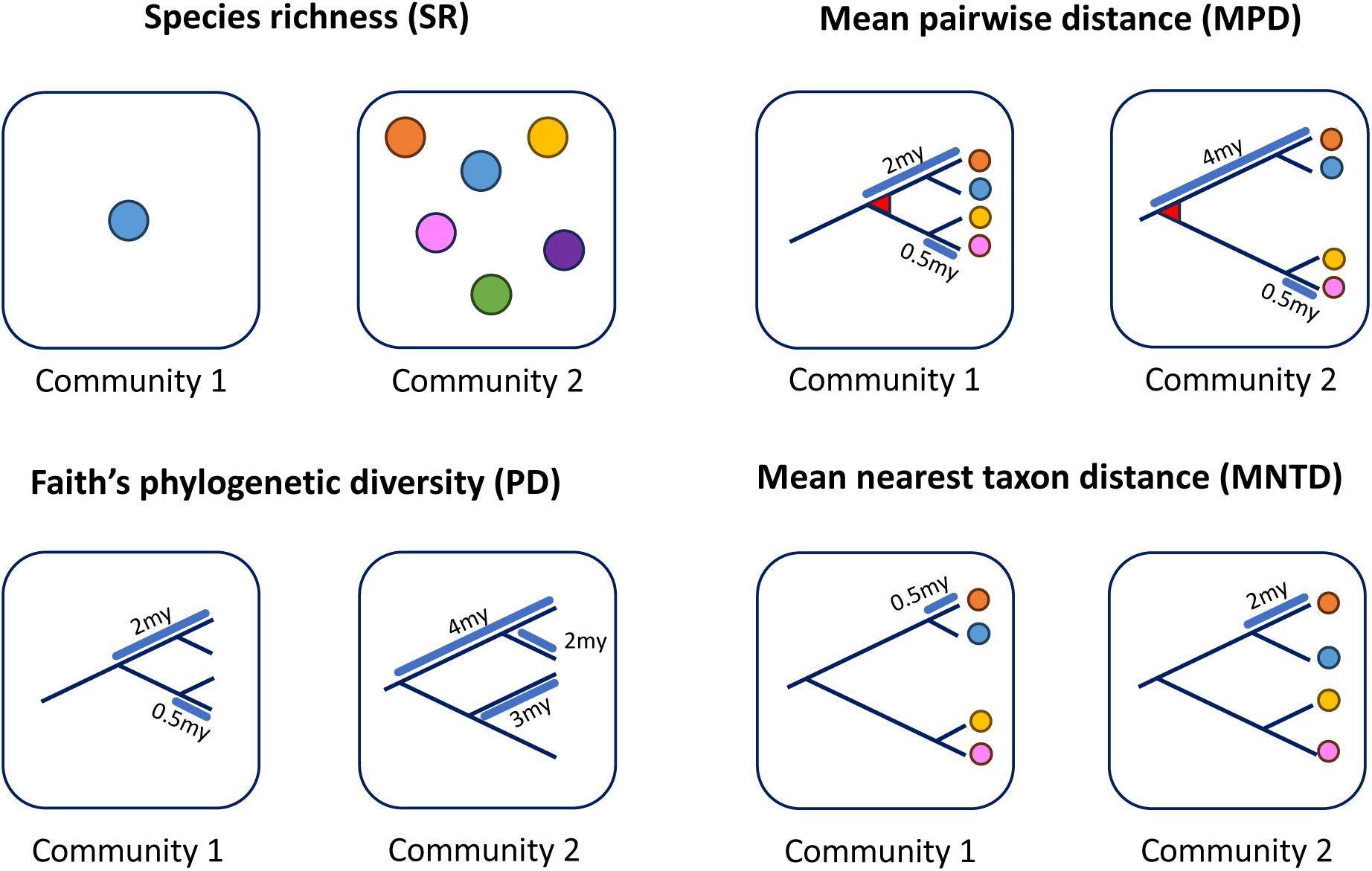
Summary of metrics used to investigate biodiversity at the Knepp Estate and Boothby Wildland. my = million years. **SR** allows comparison of simple species count. Community 1 has low SR (n = 1) compared with community 2 (n = 6). **Faith’s PD** calculates differences in relatedness. Here, the sum of branch lengths in community 1 is lower than in community 2, as community 1’s species diverged more recently. **MPD** takes the mean phylogenetic pairwise distances for each species in a community (orange to blue, orange to yellow, orange to pink, blue to orange etc.). In community 1, because of the shorter time since divergence to the node for the last common ancestor of all 4 species (marked red) than community 2, MDP would be less in community 1. **MNTD** measures mean phylogenetic distance of each species in a community to its nearest neighbour (orange to blue, blue to orange, yellow to pink, pink to yellow). Here, the most recent common ancestor for all 4 species in the community remains in the same place for both communities 1 and 2, but MNTD is impacted by the longer time since divergence for the pairs of species in community 2. For both MPD and MNTD, higher values indicate communities of species that are more distantly related, with common ancestors deeper in history. Lower values indicate relatively more clustered phylogenies, with species more closely related. Here we use Standard Effect Sizes of MPD and MNTD (SES_MPD_ or SES_MNTD_, respectively), whose values may be negative or positive. Negative SES_MPD_ or SES_MNTD_ values indicate lower phylogenetic diversity in an assemblage than would be expected by chance (phylogenetic clustering; assessed by comparison with a bootstrapped null distribution), whereas positive values indicate higher phylogenetic diversity than would be expected by chance (phylogenetic dispersion; also in comparison with a null distribution).

**SR** is a simple measure of taxonomic diversity: a basic count of the number of species in each 1m^2^ plot. **Faith’s PD** (Faith, 1992) can also be used as a measure of richness (Chao et al., 2016) but it additionally captures the amount of evolutionary history in an assemblage (Faith, 1992), revealing the influence of possible evolutionary factors on community assembly (Simpson and Pearse, 2021). To calculate Faith’s PD we used the Zanne et al. (2014) phylogeny and added missing species using pez::congeneric.merge (Pearse et al., 2015). Two metrics of phylogenetic structure were used to analyse divergence between assemblages: **SES_MPD_** and **SES_MNTD_**. These metrics allowed us to make inferences about assemblages compared with the expectation under a null model using random sampling of the local phylogeny (Webb et al., 2002, Kembel, 2009). In both cases, the standard effect size (SES) is a comparison of the observed measurement (MPD or MNTD) with its expectation under a defined null model – here a species-shuffled randomised null community (Kembel et al., 2010). In SES metrics, zero is the value for the null community: values greater than zero indicate overdispersion in a phylogeny, i.e. species that are more distantly related to one another than expected by chance. Values below zero indicate species within an assemblage that are more closely related than would be expected (Kembel, 2010, Kembel, 2009). SES_MPD_ can identify tree-wide clustering while SES_MNTD_ is more effective at identifying patterning close to the tips of trees (Kembel, 2010). These patterns can be used to infer possible ecological influences on community assembly. Assemblages of closely-related species have been described as implying habitat filtering, while collocated, distantly-related species may suggest competitive exclusion (Webb et al., 2002). This mapping of pattern onto process is based on assumptions (Cavender-Bares, 2009, Mayfield, 2010), but these biodiversity metrics are sensitive to community assembly and so they are good metrics of community structure.

We provide rank / abundance plots (Whittaker, 1960) to visualise changes in each community over time, and investigate differences by comparing the natural histories of each site’s most prevalent species in each year of the study. We also present a non-metric multi-dimensional scaling (NMDS) plot to visualise the assemblages over time (NMDS calculated with 2 dimensions; stress = 0.146). A Bray-Curtis distance matrix was used to generate the NMDS and ellipses are drawn from 95% confidence intervals estimated assuming the points are drawn from multivariate normal distributions (see ggplot2::stat_ellipse (Fox and Weisberg, 2011, Wickham, 2016). One site (Knepp M333) was removed from the data set to perform this analysis as zero species were recorded there.

### Spatial analysis

We extended our analysis by incorporating the nested spatial arrangement of our survey sites to investigate variation at different scales across these landscapes (β-diversity) (Anderson et al., 2011). We used hierarchical scaling and variance components analysis (see Crawley, 2012) to identify scale sensitivities (Swenson et al., 2006) that may be common across rewilded ex-agricultural landscapes. Different processes are thought to impact plant biodiversity at different spatial scales (Crawley and Harral, 2001), and we aimed to reveal signatures of this. Each level of the nested fractal arrangement (Figure 1 A-C) captured a different spatial scale (hierarchical scaling) from minor triads spanning 100m to full fractals spanning 900m. An additional level, block, was introduced at Knepp, where the three major parcels of land have been subject to different management strategies. Block was not introduced at Boothby, where all the land has been subject to the same management regime (crop rotation) over time. We conducted variance components analysis across each α-diversity metric using a Bayesian hierarchical model, following Simpson and Pearse (2021). This approach allowed us to reveal the proportion of variance in each biodiversity metric that was accounted for at the different spatial scales in each system. To determine the significance of our results and account for spatial autocorrelation we calculated null permutations for each metric, repeating the variance components analysis (n = 999) with each repeat conducted on a randomisation of our observed data.

We repeated the same methodology for both locations in 2022, 2023, and 2024, collecting comparable data. Our method was flexible enough to account for the fact that our case study locations were different sizes and shapes (Figure 1 D-E), and that the land at Boothby had been subject to the same treatment (decades of agricultural crop rotation) while Knepp was partitioned into three distinct blocks with different histories. By using self-similar nested fractals with a statistically conservative analysis, we could directly compare metrics across these different locations.

## Results

### Comparison of rewilded and baseline agricultural site: Knepp Estate vs Boothby Wildland, 2022

We included data for 110 sites in 2022 (32 at Knepp, 78 at Boothby) to compare community structure and composition at Knepp with a baseline at Boothby Wildland (“year 0”).

SR and Faith’s PD revealed greater diversity at Knepp than Boothby. Overall species richness was higher at Knepp (102) than Boothby (78). The median number of species found at each 1m^2^ site was 9 at Knepp (interquartile range (IQR) Knepp = 5), but just 1 at Boothby where many sites contained only one of the three monoculture crop species (IQR Boothby = 1). A similar trend was found in phylogenetic diversity – median Faith’s PD at Knepp (median = 908myr, IQR = 403myr) was more than 2x greater than at Boothby (median = 391myr, IQR = 188myr). As expected (discussed in Tucker and Cadotte (2013)), strong association was found between PD and SR in 2022 and subsequent years (all Spearman’s rho > 0.775 with p < 0.001, full details in supporting information).

Results of SES_MPD_ and SES_MNTD_ analyses were mixed. Median SES_MPD_ was positive at both Knepp (median = 0.179, IQR = 1.58) and Boothby (median = 0.170, IQR = 0.949), indicating that in both cases species were less closely related than would be expected by chance. Average SES_MNTD_ was negative at Knepp (median = −1.00, IQR = 1.25), indicating species were more closely related that would be expected by chance, but positive at Boothby (median = 0.631, IQR = 1.06).

Detail on the natural histories of the dominant species in each location adds depth to these results (Stace, 2010). The ten most-recorded species at Knepp in 2022 (listed from most-frequently observed in plots to least: *Agrostis stolonifera*, *Holcus lanatus*, *Ranunculus repens*, *Trifolium repens*, *Agrostis capillaris*, *Pulicaria dysenterica*, *Senecio jacobaea*, *Cirsium arvense*, *Phleum bertolonii*, *Quercus robur*) were native British species and each species was recorded at between 5 and 7 sites. At Boothby, the three most-recorded species were all non-native crop plants (*Triticum aestivum, Hordeum distichum,* and *Vicia faba*) and were recorded in 24, 22, and 18 plots respectively, confirming the dominance of monocultures across the landscape. Two of the ten most recorded species at Boothby were “archaeophytes”, known to be associated with human activity and to have been in the British Isles since medieval times but with uncertain deeper provenance (*Chenopodium ficifolium, Viola arvensis*), such that just five of the ten most-recorded species were native British species (*Festuca rubra, Poa trivialis, Rubus fructicosus, Phleum pratense, Ranunculus repens*).

Variance components analysis revealed greater spatial structure at Knepp than Boothby (Figure 3). Arrangement by block at Knepp (grouping into northern, middle and southern) explained the most variance in variance components analyses for SR, PD, and SES_MPD_. Further analysis of biodiversity metrics at Knepp broken down by block suggest that lower diversity in the middle block may be driving these findings (Table 1). At Boothby, where spatial structure was identified we did not identify any recurring patterns.

**Figure 3.**
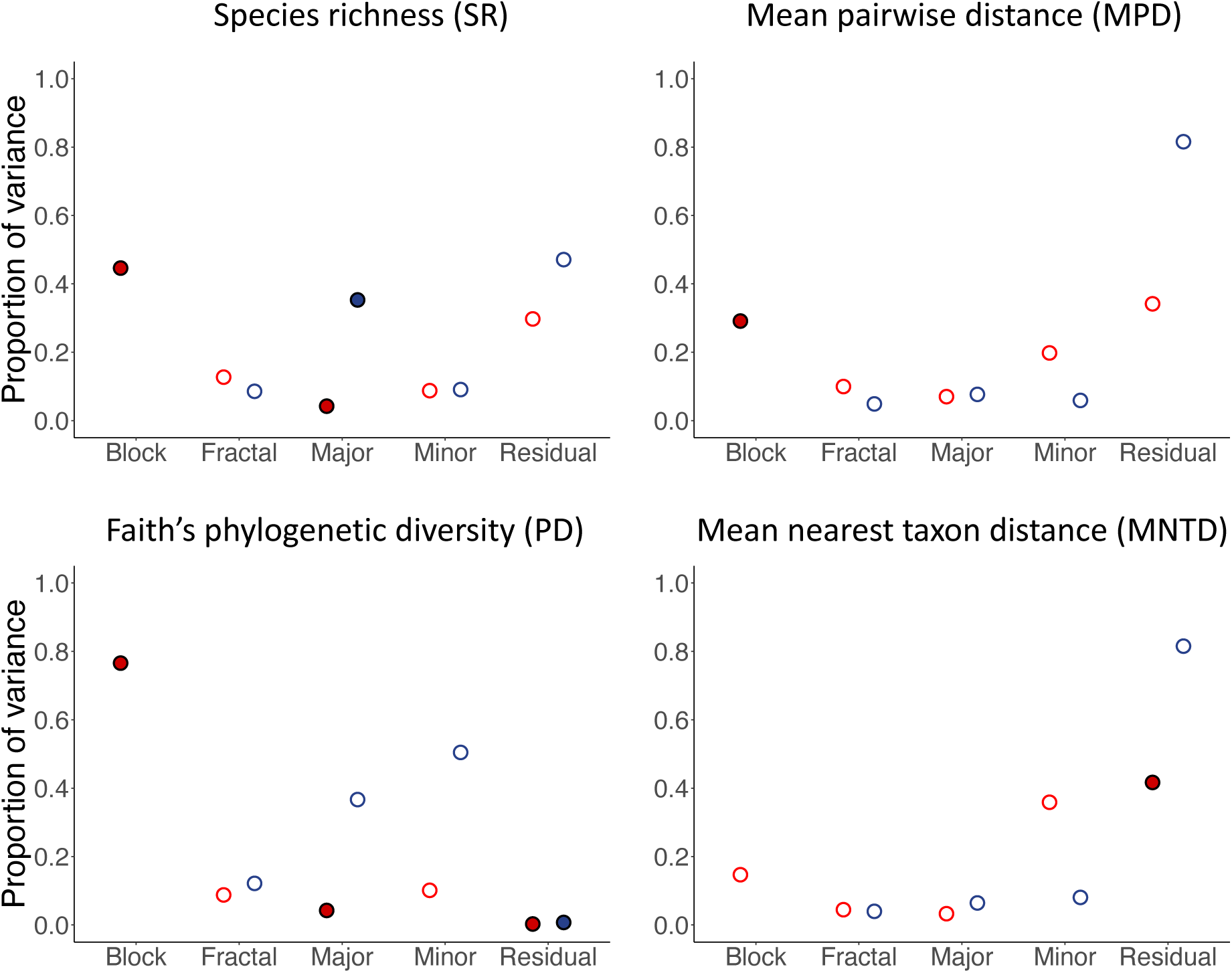
Variance components analysis outputs for four α-diversity metrics in 2022, showing contrasts in β-diversity between a rewilded system (the Knepp Estate – results in red) and a baseline system (Boothby Wildland – results in blue). Horizontal axes show the hierarchical spatial groupings - an additional level (block) was introduced at Knepp to account for the different land management strategies that have been implemented across the estate (northern, middle, and southern blocks). Vertical axes show the proportion of variance explained by landscape. A large proportion of variance at residual level indicates lack of spatial structure, i.e. variance in a metric cannot be explained by the hierarchical spatial design of the survey. Where variance in metrics was explained by the landscape, different signatures in variance portioning are revealed. Circles indicate the observed metric proportion of variance. For simplicity of presentation results are shown as either significant or not significant; full null distributions are shown in supporting information. Significant results are represented by filled circles (red for Knepp, blue for Boothby). p-values of results that are significantly different from the mean of the null permutation distribution: **SR** Knepp (block = 0.002, major triad = 0.04), **SR** Boothby (major triad = 0.028). **Faith’s PD** Knepp (block = 0.002, major triad = 0.008, residual = 0.024), **Faith’s PD** Boothby (residual = 0.024). **SES_MPD_** Knepp (block = 0.012). **SES_MNTD_** Knepp (residual = 0.032).

**Table 1.** Grazing regimes in different blocks of the Knepp Estate drive biodiversity differences (2022 data). The northern and southern blocks are broadly similar, but the middle block has a very different character with roughly half the median SR and PD of the other blocks. The standardised effect size of mean pairwise distance (SES_MPD_) shows phylogenetically clustered assemblages in the middle block (negative median SES_MPD_) compared with more phylogenetically dispersed assemblages in the northern and southern blocks. These results, driven by differences in landscape management approaches, underpin the influence of grouping by block in variance components analysis. Species richness (SR) to nearest whole number of species, other figures to 3 significant figures; PD reported in units of millions of years and other metrics are dimensionless test-statistics.

|  | SR |  | PD |  | SES<sub>MPD</sub> |  | SES<sub>MNTD</sub> |  |
| --- | --- | --- | --- | --- | --- | --- | --- | --- |
| Block | Median | IQR | Median | IQR | Median | IQR | Median | IQR |
| Northern | 10 | 5 | 1030 | 319 | 0.639 | 0.710 | -1.13 | 1.84 |
| Middle | 5 | 2 | 508 | 224 | -0.817 | 1.61 | -0.821 | 1.23 |
| Southern | 10 | 5 | 950 | 185 | 0.193 | 1.30 | -0.566 | 1.61 |

### Monitoring changing community compositions in early rewilding (2022-2024)

We repeated our analyses for each year of fieldwork data (2022, 2023, 2024) to monitor changes in these two systems over time. α−diversity metrics at Boothby changed dramatically over the 3 years captured by this study. SR more than doubled (SR 2022 = 78, median = 1, IQR = 1. SR 2024 = 186, median = 9, IQR = 5), as did Faith’s PD (PD 2022 median = 391, IQR = 188. PD 2024 median = 875, IQR = 310). In 2024, total SR at Boothby was greater than that recorded at Knepp (SR 2024 Knepp = 162, Boothby = 182). Both median SES_MPD_ and median SES_MNTD_ became considerably more negative at Boothby between 2022 and 2024 (SES_MPD_ 2022 median = 0.170, IQR = 0.949. SES_MPD_ 2024 median = −0.569, IQR = 0.513. SES_MNTD_ 2022 median = 0.631, IQR = 1.06. SES_MNTD_ 2024 median = −1.465, IQR = 1.40) suggesting that one impact of rewilding is the formation of communities that are more closely related than would be expected by chance (i.e. phylogenetic clustering).

Rank / abundance plots and NMDS visualise these changes (Figure 4; Figure 5) and qualitative comparison of the sites with reference to species’ natural histories (Stace, 2010) provides more detail. From 2022 to 2024 a decline in the number and dominance of non-native crop species was charted, with increasing numbers of records of native British species (full details supporting information).

**Figure 4.**
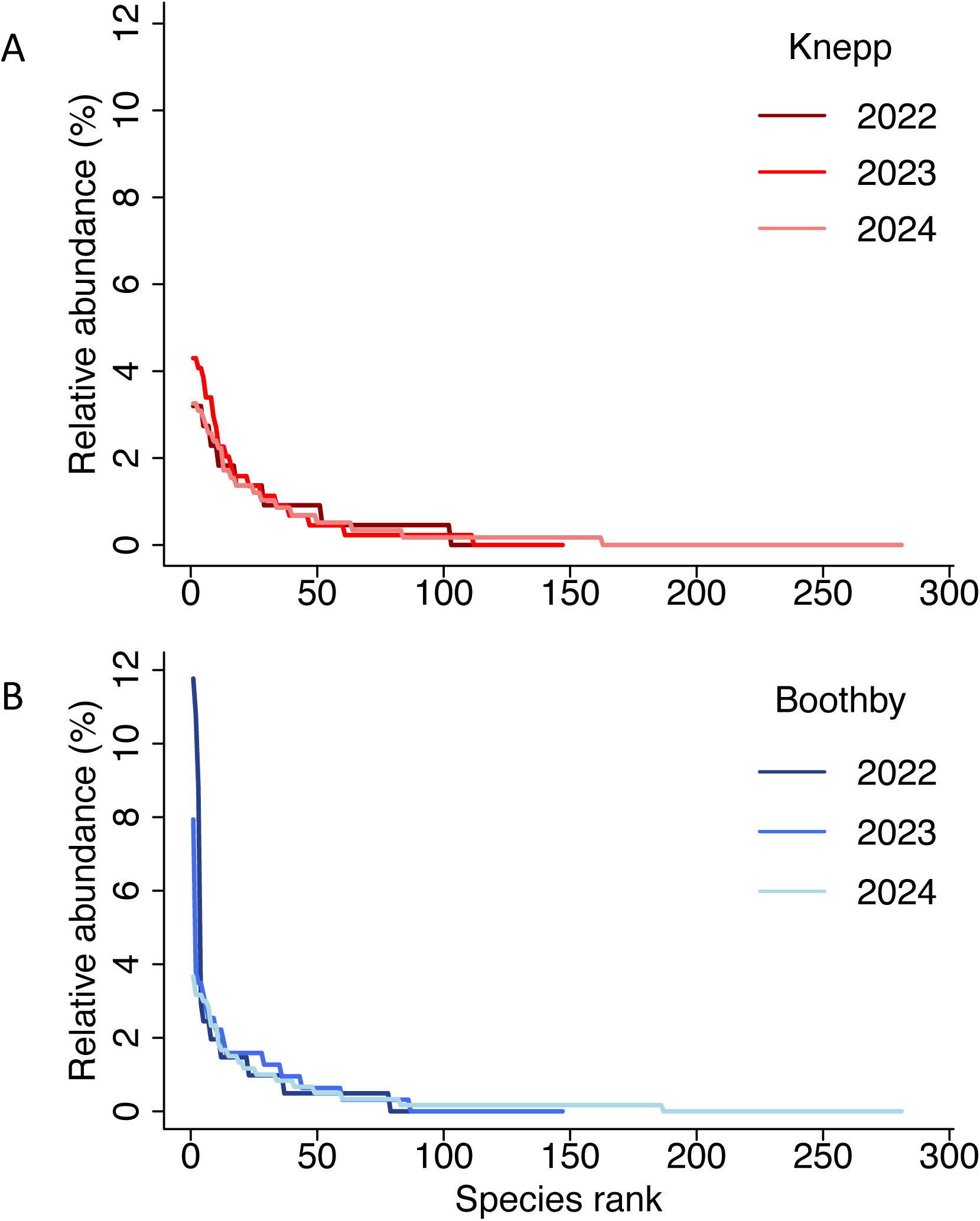
Rank-abundance plots showing changes in community composition and structure from 2022 to 2024. Horizontal axes show species sorted into descending order of abundance such that species rank = 1 is the species most commonly recorded in survey data. Vertical axes show relative abundance of each ranked species as a percentage. **A.** Limited changes through time at the Knepp Estate between 2022 and 2024. The species with highest relative abundance is similar each year (2022 = 3.20%, 2023 = 4.30%, 2024 = 3.26%), and the shape of the three curves nearly identical. This suggests that the plant community at Knepp was relatively stable during this time. **B.** Data for Boothby Wildland, on the other hand, reveals a rapidly changing community with declining dominance of crop species. The number 1 ranked species accounts for much greater relative abundance in Boothby than in Knepp in 2022 (11.8%) and in 2023 (7.94%). By 2024 the curve has come to resemble that of Knepp, with the number one ranked species accounting for a similar relative abundance (3.67%). This represents the declining dominance of crop species and commensurate increasing plant diversity and evenness over time.

**Figure 5.**
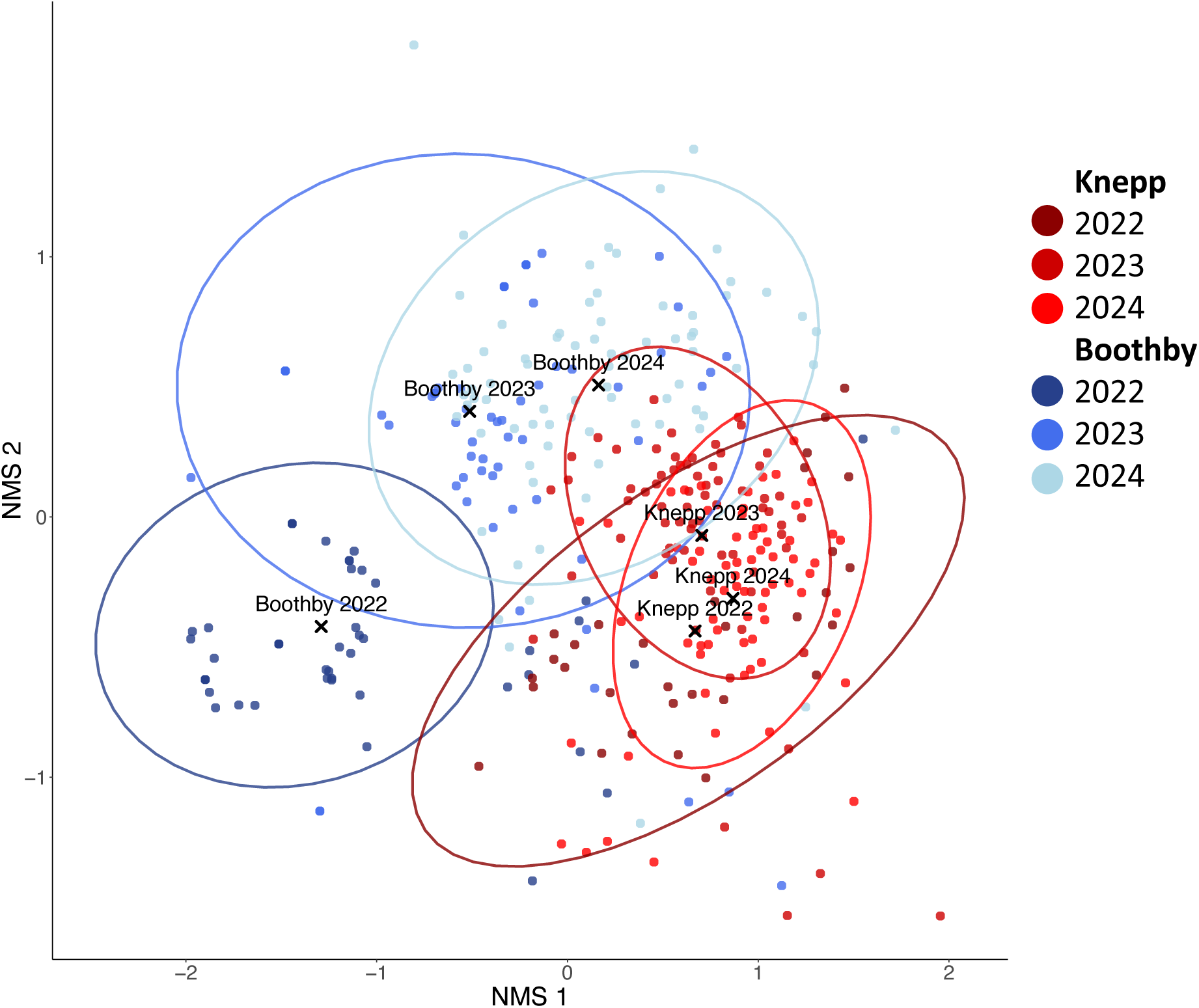
Non-metric multi-dimensional scaling (NMDS) suggests biodiversity at Boothby Wildland (blue) is moving to more closely resemble that at the Knepp Estate (red). Ellipses are shown for each site in each year (level = 95%). Separation of centroids from ellipses of other years suggests significant differences. The three red ellipses for Knepp are clustered together, and all three centroids within all three ellipses. The blue ellipses for Boothby are more dispersed in the NMDS space. Boothby 2022 is distinct from both Boothby 2023 and 2024. This supports the argument that the plant community at Knepp remained relatively stable during this study period, while that at Boothby was changing year-on-year.

Analysis of spatial structure at Knepp and Boothby yielded mixed results, but does support the idea that community structure was relatively stable at Knepp while changing at Boothby. The clearest trends emerge in the results for Knepp. In all three years, significant results were recorded for the large proportion of variance in PD explained by grouping at block level and the small amount explained by grouping at major and residual levels. The small effect of the fractal and major grouping categories was also a consistent trend at Knepp. Results for Boothby, on the other hand, returned fewer significant results and no trends were consistent across all three years. Most significant results at Boothby derived from the 2023 analysis, but none were returned in 2024. Results for SES_MPD_ and SES_MNTD_ show that very small proportions of variance were explained by landscape at Boothby in all three years (almost all variance assigned to residual), but at Knepp some structure was identified at the block and minor triad levels. Results for all metrics at Knepp and Boothby across the three years are presented in Figure 6.

**Figure 6.**
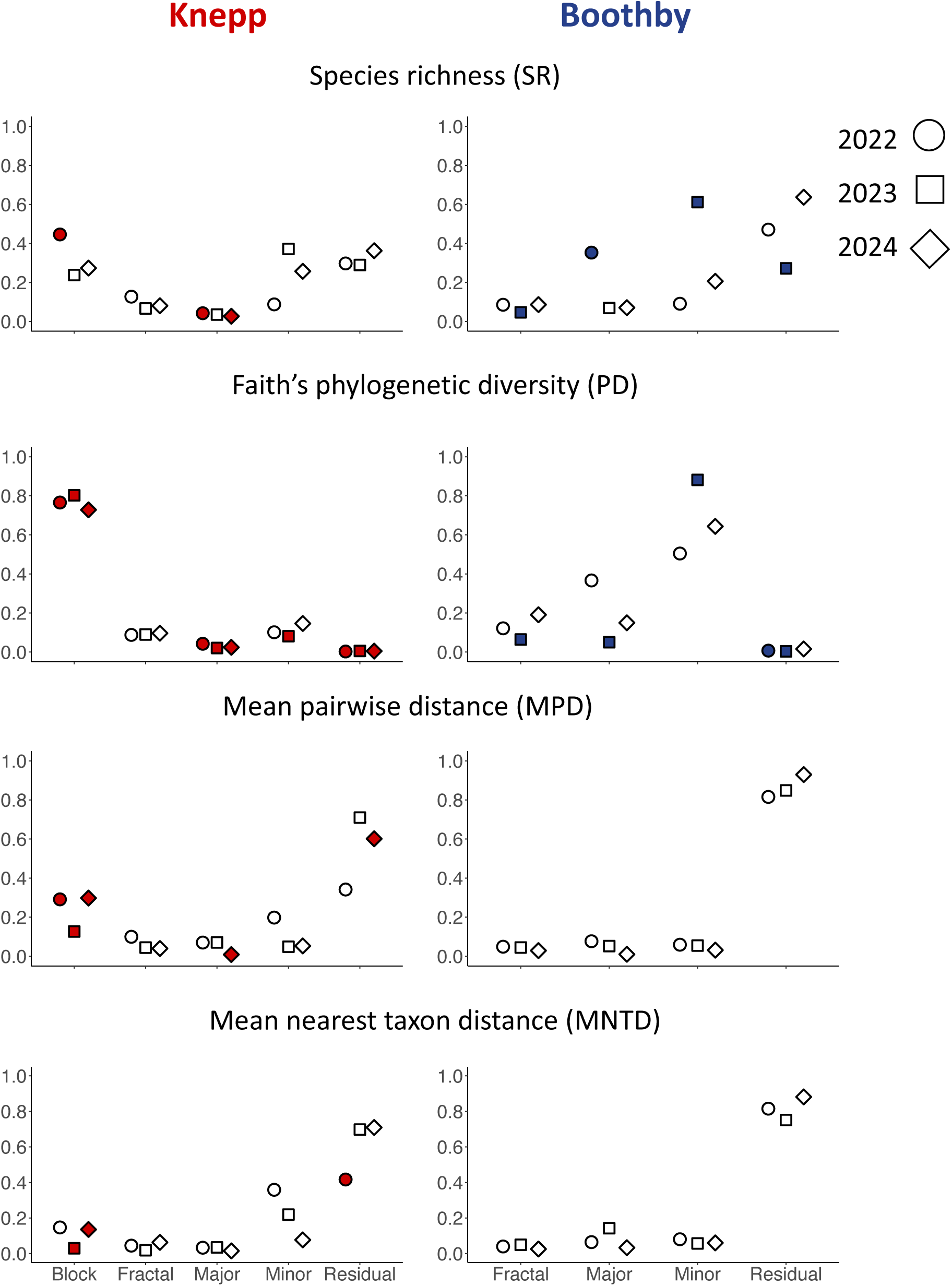
Overall results of variance components analysis for four α-diversity metrics over 3 years, showing contrasts in β-diversity between a rewilded system (the Knepp Estate) and a baseline system (Boothby Wildland). x axes show the hierarchical spatial groupings – including block level, introduced at Knepp to account for the different land management strategies. y axes show the proportion of variance explained by landscape. The left-hand plots show data from Knepp and right-hand from Boothby. Plotted shapes indicate the observed metric proportion of variance in this study. Circles: 2022, squares: 2023, diamonds: 2024. Significant results are represented by filled shapes: red for Knepp, blue for Boothby. See supporting information for full results for variance components analyses for all metrics.

## Discussion

We studied the structure of biodiversity in one agricultural landscape (Boothby Wildland) and another >20 years into rewilding (the Knepp Estate). We use this case study to describe quantitatively the way in which land reclaimed from an agricultural baseline may be expected to change in community structure over time by comparing α− and β-diversity metrics between the two locations. Our study included three years of data collection, including the first two years of transition for Boothby from an agricultural to rewilded landscape. Based on our findings, we make some inferences about rewilding strategies, particularly the importance of managed herbivory, and we also provide recommendations for monitoring and evaluation of rewilding projects. We show that, in our sites, the trajectory towards recovery from a degraded agricultural state is characterised by rapid change and that recovery can be detected first at small spatial scales. We argue that spatial metrics for SES_MPD_ and SES_MNTD_ may be a key metric identifier for distinguishing between rewilded and baseline agricultural landscapes more generally.

### Comparison of rewilded and baseline agricultural site: Knepp Estate vs Boothby Wildland, 2022

Our results revealed differences in structure between the mature system and the baseline landscape at year 0. We confirmed expectations of limited α− and β−diversity in the agricultural landscape at Boothby, finding greater diversity and increased spatial structure in the rewilded landscape at Knepp. A key trend at Knepp was that, in variance composition analysis (Figure 3), large proportions of variance were explained by grouping at block level. From this we infer that managed herbivory is a major factor determining the structure of biodiversity in rewilded landscapes.

Herbivory is a key element in rewilding (Gordon et al., 2021) and reintroductions of large herbivores have been found to lead to improved connectivity in fragmented landscapes (Berti et al., 2020). However, herbivore population densities require careful management in the absence of predator populations (Cooke, 2021, Fattorini et al., 2020), and this will be a significant consideration for any UK project. We were able to explain most variation at Knepp by grouping at block level (northern, middle, and southern), where different herbivory management regimes have been implemented in the years since farming was halted (Kirby, 2020). There is no analogous grouping level at Boothby, where the land has been subject to the same treatment for many years. The middle block at Knepp had a very different diversity profile compared with the other two blocks (Table 1), and we argue this was due in large part to its different, more intensive, grazing regime. The number of grazing animals (deer and cattle) has been kept intentionally high in the middle block to maintain a parkland appearance (Tree, 2018), favouring grasses and herbaceous flowering species rather than woody shrubs and brambles (Putman and Moore, 1998). High grazing pressure leads to less diverse assemblages (Reed et al., 2022) and the difference between this regime and those in the northern and southern blocks likely explains the block-level effects observed. At Boothby, where herbivores have not yet been kept as part of the management strategy, most variance was captured at residual level in three out of four variance components analyses (all except Faith’s PD) indicating a lack of spatial structure (Figure 3). α-diversity metrics also revealed homogenisation across the landscape (see supporting information). The fields have been subject to crop rotation and intensive agricultural processes for decades and, in 2022, 86% of the total site area was given over to the growth of one of three monocultures (pers. obs. Ivan de Klee). Many natural processes of community assembly (e.g., dispersal, habitat filtering, and competitive exclusion) have therefore been disrupted as the dominant crop species were grown from purchased and planted seed rather than naturally dispersed populations, and non-crop species were actively removed by weeding.

Our results confirm that herbivory management is of great importance to rewilding outcomes. Both heavy grazing (Knepp, middle block) and intensive agricultural practices (Boothby) can lead to decreased biodiversity by reducing landscape heterogeneity. Grazing by herbivores is thought to play a significant role in the growth and reproduction of plants (Coughenour, 1985, Forrestel et al., 2017), with the presence of large herbivores attributed to increased richness and diversity of plant communities (Collins and Calabrese, 2012, Koerner et al., 2014). Heavy grazing, however, can lead to biotic homogenisation (Salgado-Luarte et al., 2019) and can degrade soil sufficiently to threaten its subsequent recovery by excluding the return of species lost by overgrazing (Villamil et al., 2001). Thus, only groups that can adapt to the new conditions are able to persevere. At Knepp, selective grazing of major groups of species from much of the middle block would be expected to limit both SR and Faith’s PD. Species would not be grazed (and so filtered) at random, and we would expect to see the evidence of this structuring across the Knepp Estate. Indeed, block explained the majority of variance in three out of four metrics studied at Knepp (Figure 3), and further investigation into the diversity of the northern, middle, and southern blocks (Table 1) confirmed the reduced diversity of the middle block. This demonstrates an important aspect of our methodology: pooling the diversity at Knepp would mask the evidence that the middle block contained a distinct and quite different community, a fact that was revealed by variance components analysis.

### Monitoring changing community compositions through rewilding (2022-2024)

We wanted to evaluate whether diversity at Boothby would change during the study period in a way consistent with a trajectory towards the structure of Knepp. We show that the early years of transition from a baseline agricultural state are characterised by rapid change and that spatial signatures of recovery can first be detected at small scale. These changes are being driven by dramatically increasing species richness and it is likely that recruitment from the seedbank and local dispersal are responsible for much of the increased diversity, rather than vegetative growth. We argue that two biodiversity metrics, SES_MPD_ and SES_MNTD_, may be key indicator metrics for measuring recovery progress. The plant community at the Knepp Estate was relatively stable from 2022-2024 compared to that of Boothby Wildland. We observed changing plant community composition at Boothby, with increases in diversity driven by a decline in dominance of non-native crop species and increasing prevalence of native British species (Figure 4, also supporting information). We suggest that the landscape at Boothby may be shifting to more closely resemble that of Knepp (Figure 5). Our spatial analyses support this to some extent. Results of variance components analysis at Knepp were consistent between 2022 and 2024 (Figure 6) with the observed proportion of variance for each metric in each year mostly tightly grouped. At Boothby, on the other hand, results changed each year for species richness and Faith’s PD while remaining unchanged for SES_MPD_ and SES_MNTD_ (with a negligible amount of variance explained by landscape).

These results inform two arguments. Firstly, where structure was evident at Boothby it explained most variance at minor triad level (Figure 6) suggesting that in the early stages of landscape recovery spatial metrics can detect changes first at the smaller scales. This is in contrast with the more mature system at Knepp where the largest spatial scale (block) tended to capture the most variance. This is likely linked to the transient dynamics present in the recovering landscape of Boothby, where high disturbance conditions allow for ingress of pioneer (ruderal) species compared with the more established dynamics at Knepp following more than 2 decades of rewilding, where broader site conditions – for example soil type and herbivory – contribute more to the composition of the plant community (Grime 1974; Grime 1977). Secondly, spatial metrics for SES_MPD_ and SES_MNTD_ best distinguished between the degraded and rewilded landscapes we studied. These metrics consistently revealed some structure in the rewilded landscape at Knepp but almost none in the baseline landscape at Boothby. These metrics may, therefore, represent a key identifier for evaluating landscape recovery through rewilding and further research should be focused on investigating this further.

## Conclusions

The Lawton Report (Lawton et al., 2010), a highly influential review of England’s wildlife sites (Rose et al., 2018), established that England required “more, bigger, better and joined” ecological networks (Lawton et al., 2010). The UK has recently committed to a 25-year plan to restore landscapes and protect biodiversity (Defra, 2018) and more land is due to be turned over to rewilding in the coming years (The Wildlife Trusts, 2023, Burford, 2023). While this is positive, we still do not know what the trajectory of such sites should look like, or what the long-term biodiversity impacts will be. If rewilding continues to grow in popularity, increasing numbers of small rewilded spaces will be introduced across the UK. Since 70% of the UK’s land area is agricultural, rewilded spaces will often begin from this baseline. It is therefore critical that we understand how rewilding can shape biodiversity in these habitats (Tscharntke et al., 2005). Developing this understanding will require improved methods for monitoring. A lack of monitoring has been highlighted as a key evidence gap in rewilding projects in the UK (Bailey et al, 2022). Monitoring methodologies will need to be simple and cheap to implement, and this study demonstrates the effectiveness of our method. This survey design can be applied across terrestrial landscapes of varying sizes and shapes to create directly comparable results with relatively low sampling effort, and it can be flexibly adapted (e.g. to capture the management of different parcels of land, as at the Knepp Estate). We also showed that in landscapes that are known to be homogenous, surveying can be simplified to maximise the number of sites that can be included if surveying time is restricted, allowing fieldwork to be organised and targeted efficiently, within realistic constraints of time and money available (Marsh and Ewers, 2013).

This case study establishes that spatial variation is a valuable measure to complement consideration of α−diversity metrics when assessing landscape recovery through rewilding. We show that different landscape management strategies can impact the development of community structure, particularly the inclusion and population management of herbivore species. Our methodology may provide the first stage in identifying a direction of community change that could be generalisable to any agricultural landscape recovery project in the UK. To validate this, it would be valuable to observe changes at Boothby over the coming years, where we expect partitioning of variance will come to further resemble that at Knepp. It will be useful to know both if, and how quickly, this occurs to inform monitoring strategies for future rewilding projects taking landscapes from an agricultural baseline. This survey design should also be repeated on as many agricultural baselines and rewilded systems as possible and the results pooled to build a useful picture of general trends. This could be used to inform guidelines on monitoring and evaluating progress in rewilding in a quantitative way as the UK continues to build towards more, better, more natural spaces.

