## Supplement for "Using landscape biodiversity metrics to assess rewilding: a space-for-time comparison between the Knepp Estate and an agricultural baseline at Boothby Wildland"

**Supporting Information**

Supporting information is broken down into sections containing:

1. Additional detail on the extended methodology employed at Boothby Wildland in 2022 and 2023 to maximise the number of data points that could be included in analyses.
2. Full details of the natural histories of the most prevalent species recorded in each site in each of the 3 survey years.
3. α-diversity metrics and the results of variance components analysis in each year of the study (2022, 2023, and 2024) including null distributions.
4. Results of correlation tests between species richness and Faith’s phylogenetic diversity

**Section 1**

Section 1 provides additional detail on the extended methodology employed at Boothby Wildland in 2022 and 2023 to maximise the number of data points that could be included in analyses. This approach allowed field work time to be focussed on more complex sites with larger numbers of species where large numbers of sites were within fields of monoculture crops.

*2022*

Boothby was still an active farm in 2022, so much of its area was dominated by cash crops (wheat, barley and broad beans). We included data for 78 sites in analyses. 37 of these sites were surveyed using the full methodology outlined in the main text. 27 sites were assumed to be monocultures as they fell within fields where multiple full surveys had been completed. A subset of these 27 (n = 13) were photographed to confirm that only the expected crop species were present.

*2023*

In 2023, approximately one third of the total area at Boothby had been removed from farming. Many sites remained in crop fields, however, 86 sites were included in analyses for 2023. Of these, 25 were only photographed to confirm that only crop species were present.

2024

In 2024, when approximately another third of the land at Boothby had been removed from farming, the extended methodology was not employed and all sites were surveyed in full. Table S1.1 contains a breakdown of the number of sites surveyed using the extended methods in each year of the study.

**Table S1.1 Breakdown of sites surveyed in full, surveyed by photograph, and assumed to be crop monocultures.**

|  | **2022** | **2023** | **2024** |
| --- | --- | --- | --- |
| **Boothby (total sites)** | 78 | 86 | 77 |
| **Boothby (photographed)** | 14 | 25 | 0 |
| **Boothby (assumed)** | 27 | 0 | 0 |

To validate this methodology, we repeated our analyses on subsets of the data from Boothby in 2022 and 2023 (Figure S1.1, Figure S1.2). Results were near-identical in both cases, supporting the use of this flexible methodology for landscapes that are known to be homogenous. The full data set and code to reproduce statistical analyses are provided online (DOI: 10.5281/zenodo.14035907).

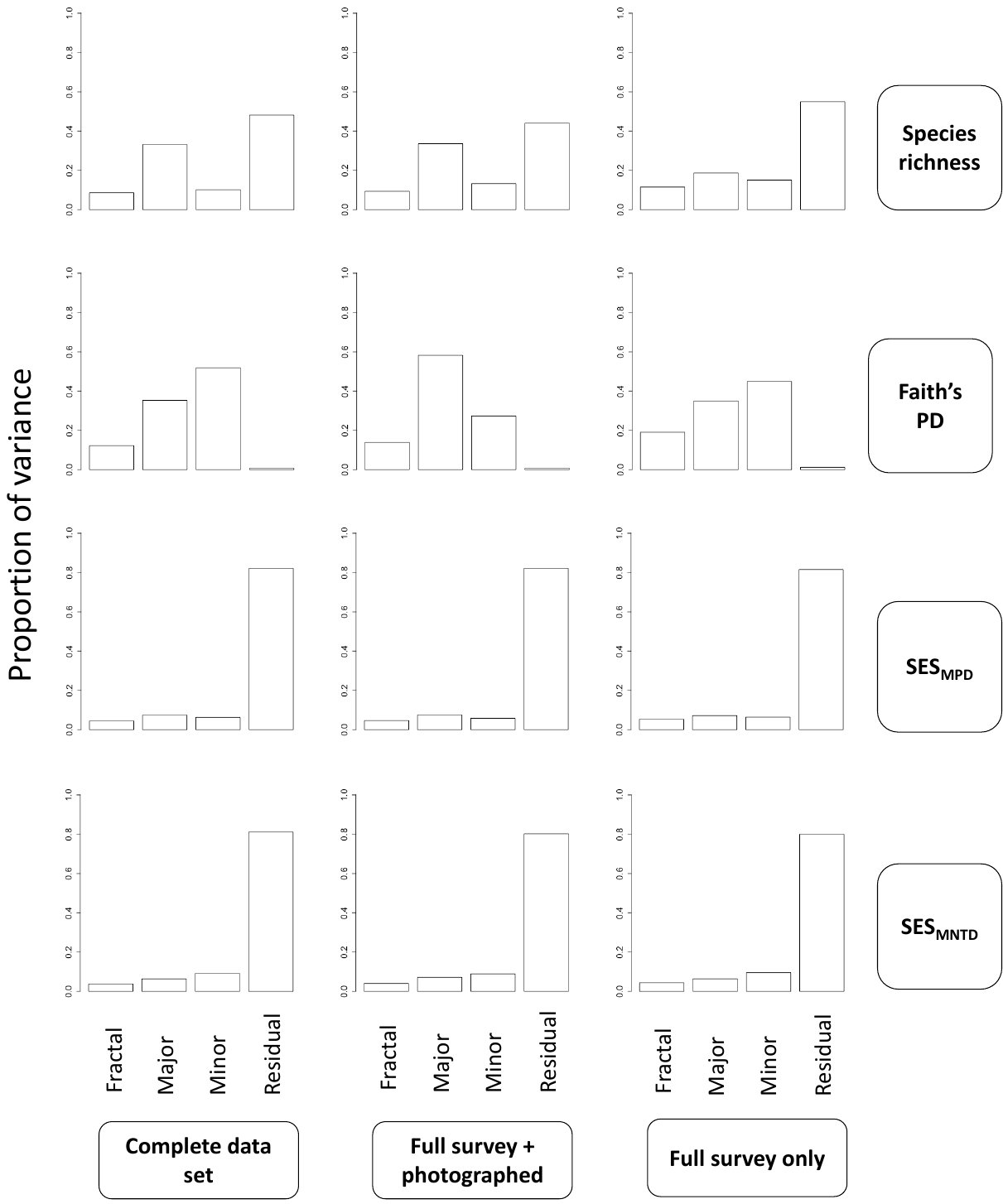

**Figure S1.1 Outputs of repeat analyses of Boothby data in 2022 to confirm extended methodology employed to maximise the number of site plant surveys included.** Left column = results of variance components analysis with all sites included (full survey methodology, photographed sites, and sites assumed to be monocultures), n = 78. Middle column = data set with assumed sites removed (sites where a full survey was conducted, and sites visited to confirm monocultures were included), n = 64. Right column = assumed and photographed sites removed (only sites where full survey methodology was conducted were included), n = 37. Qualitatively identical patterns were revealed by 14 out of 15 analyses, demonstrating the utility of maximising the range of the survey in this way where landscapes are very homogenous.

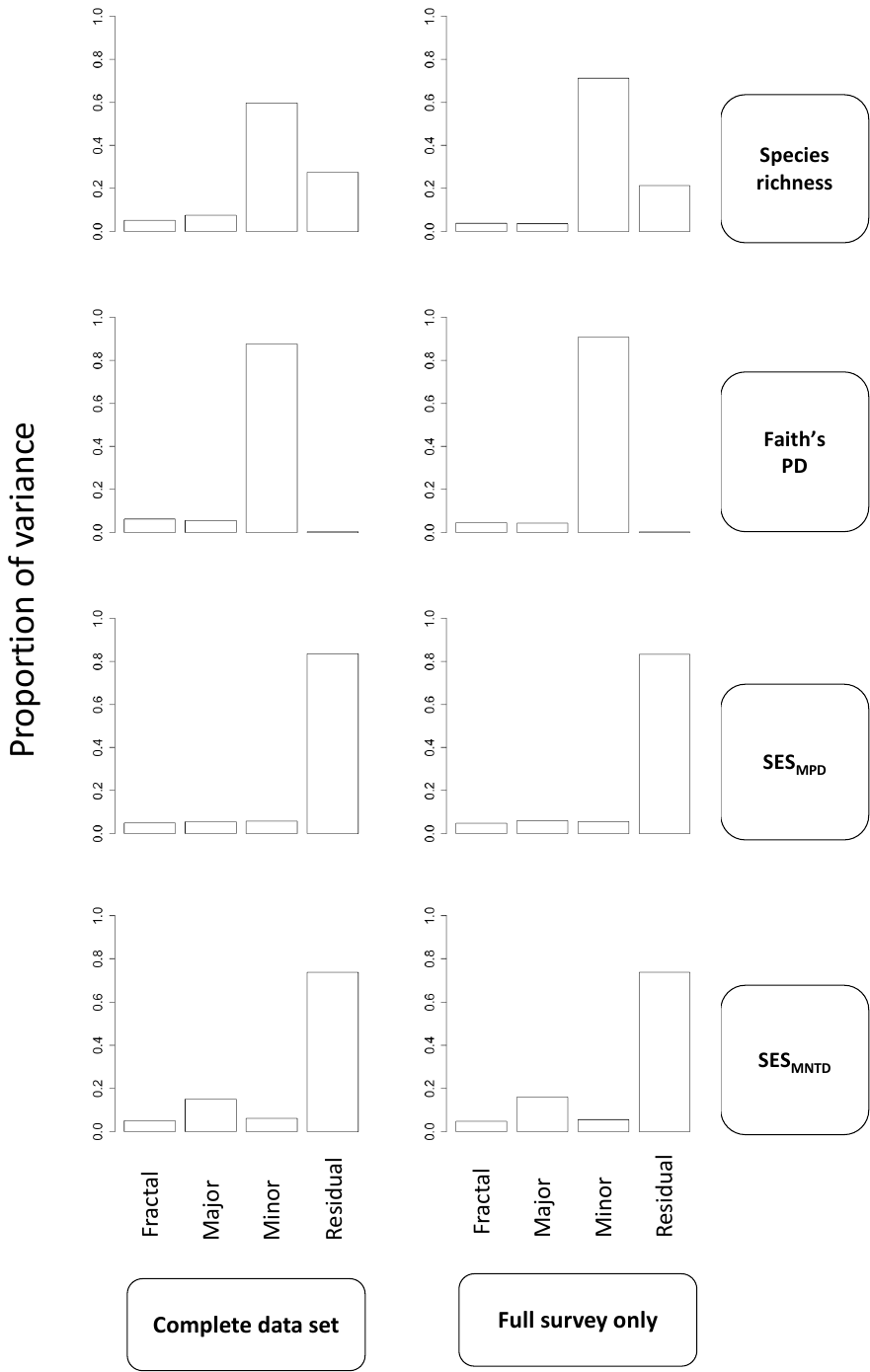

**Figure S1.2 Outputs of repeat analyses at Boothby data from 2023 to confirm extended methodology employed to maximise the number of site plant surveys included.** Left column = variance components analysis with all sites included (full survey methodology and photographed sites), n = 86. Right column = photographed sites removed (only sites where a full survey was conducted included), n = 61. Qualitatively identical patterns were revealed in all analyses, again demonstrating the utility of maximising the range of the survey in this way where landscapes are very homogenous.

**Section 2**

Section 2 provides full details of the natural histories of the most prevalent species recorded in each site in each of the 3 survey years.

**Table S2.1 List of the most-recorded species at Knepp and Boothby in 2022, 2023, and 2024.** Including number of sites in which they were recorded (#) and a record of their status as native or non-native British plants (Stace, 2010).

| **Knepp** | **#** | **Status** | **Boothby** | **#** | **Status** |
| --- | --- | --- | --- | --- | --- |
| **2022** | | | | | |
| *Agrostis stolonifera* | 7 | Native | *Triticum aestivum* | 24 | Non-native |
| *Holcus lanatus* | 7 | Native | *Hordeum distichum* | 22 | Non-native |
| *Ranunculus repens* | 7 | Native | *Vicia faba* | 18 | Non-native |
| *Trifolium repens* | 7 | Native | *Festuca rubra* | 6 | Native |
| *Agrostis capillaris* | 6 | Native | *Chenopodium ficifolium* | 5 | Archaeophyte |
| *Pulicaria dysenterica* | 6 | Native | *Poa trivialis* | 5 | Native |
| *Senecio jacobaea* | 6 | Native | *Rubus fructicosus* | 5 | Native |
| *Cirsium arvense* | 5 | Native | *Cynosurus cristatus* | 4 | Native |
| *Phleum bertolonii* | 5 | Native | *Phleum pratense* | 4 | Native |
| *Quercus robur* | 5 | Native | *Ranunculus repens* | 4 | Native |
| **2023** | | | | | |
| *Agrostis stolonifera* | 19 | Native | *Triticum aestivum* | 25 | Non-native |
| *Festuca rubra* | 19 | Native | *Vicia faba* | 12 | Non-native |
| *Holcus lanatus* | 18 | Native | *Brassica napus* | 11 | Non-native |
| *Trifolium repens* | 18 | Native | *Matricaria discoidea* | 11 | Non-native |
| *Poa trivialis* | 17 | Native | *Anthriscus sylvestris* | 10 | Native |
| *Poa pratensis* | 15 | Native | *Epilobium angustifolium* | 9 | Native |
| *Pulicaria dysenterica* | 15 | Native | *Matricaria chamomilla* | 8 | Archaeophyte |
| *Rumex crispus* | 15 | Native | *Phleum pratense* | 8 | Native |
| *Ranunculus repens* | 13 | Native | *Pulicaria dysenterica* | 8 | Native |
| *Geranium dissectum* | 12 | Archaeophyte | *Bromus sterilis* | 7 | Archaeophyte |
| **2024** | | | | | |
| *Agrostis stolonifera* | 19 | Native | *Epilobium hirsutum* | 22 | Native |
| *Poa trivialis* | 19 | Native | *Epilobium tetragonum* | 19 | Native |
| *Holcus lanatus* | 18 | Native | *Lolium perenne* | 19 | Native |
| *Trifolium repens* | 18 | Native | *Triticum aestivum* | 19 | Non-native |
| *Festuca rubra* | 17 | Native | *Bromus hordeaceus* | 18 | Native |
| *Jacobaea vulgaris* | 16 | Native | *Epilobium parviflorum* | 18 | Native |
| *Bromus hordeaceus* | 15 | Native | *Holcus lanatus* | 17 | Native |
| *Ranunculus repens* | 15 | Native | *Alopecurus myosuroides* | 14 | Archaeophyte |
| *Geranium dissectum* | 14 | Archaeophyte | *Poa pratensis* | 14 | Native |
| *Rubus fruticosus* | 14 | Native | *Sonchus asper* | 14 | Native |

**Section 3**

Section 3 contains a table of α-diversity metrics and the results of variance components analysis in each year of the study (2022, 2023, and 2024).

**Table S3.1 Results (to 3sf) of α-diversity investigation into the four metrics of study** (species richness, Faith’s PD, SES_MPD_, and SES_MNTD_) in two study locations (the Knepp Estate and Boothby Wildland), in three years of study (2022-2024). Sites = total number of 1m^2^ sites included in the analysis. myr = million years.

|  | | SR | | | PD | | MPD | | MNTD | |
| --- | --- | --- | --- | --- | --- | --- | --- | --- | --- | --- |
| Knepp | | | | | | | | | | |
| Year | Sites | Total | Median | IQR | Median  (myr) | IQR  (myr) | Median | IQR | Median | IQR |
| 2022 | 32 | 102 | 9 | 5 | 908 | 403 | 0.179 | 1.58 | -1.00 | 1.25 |
| 2023 | 70 | 111 | 9 | 6 | 962 | 402 | -0.561 | 0.745 | -1.23 | 1.09 |
| 2024 | 71 | 162 | 11 | 6 | 1140 | 502 | -0.374 | 1.04 | -1.29 | 1.14 |
| Boothby | | | | | | | | | | |
| 2022 | 78 | 78 | 1 | 1 | 391 | 188 | 0.170 | 0.949 | 0.631 | 1.06 |
| 2023 | 86 | 86 | 2 | 5 | 192 | 414 | -0.513 | 2.26 | -0.474 | 1.64 |
| 2024 | 77 | 186 | 9 | 5 | 875 | 310 | -0.569 | 0.513 | -1.47 | 1.40 |

In each figure below, x axes show the hierarchical spatial groupings. An additional level (block) was introduced at Knepp to account for the different land management strategies that have been implemented across the estate (northern, middle, and southern blocks). y axes show the proportion of variance explained by landscape. Grey violin plots show the distribution of 999 null permutations calculated for each metric and circles indicate the observed metric proportion of variance in this study. Significant results are represented by solid red circles.

*2022 variance components analysis*

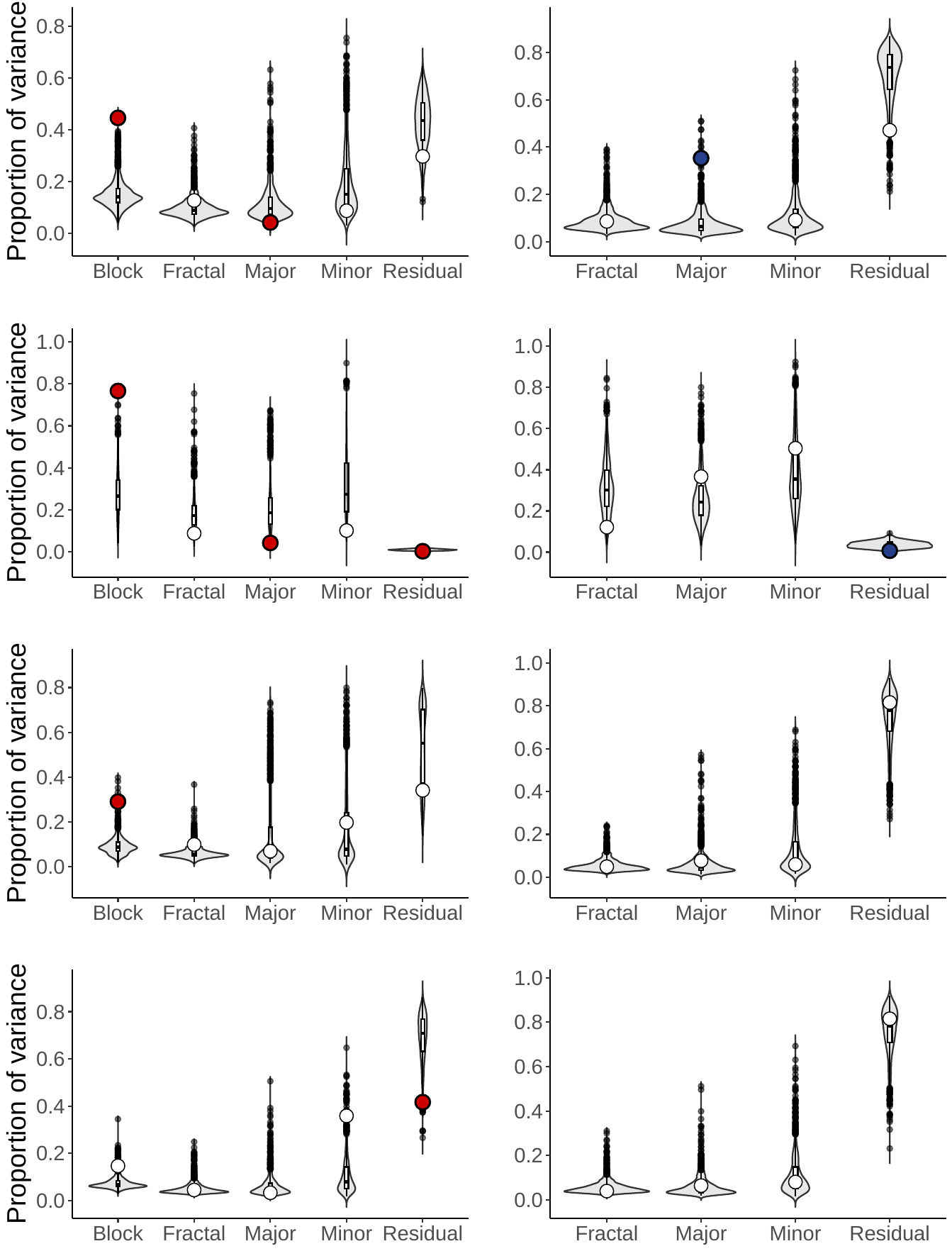

**Knepp**

**Boothby**

**Species Richness**

**Faith’s PD**

**SES_MPD_**

**SES_MNTD_**

**Figure S3.1 Pairs of variance components analysis outputs for four α-diversity metrics in 2022, showing contrasts in β-diversity between a rewilded system (the Knepp Estate) and a baseline system (Boothby Wildland).** p-values of results that are significantly different from the mean of the null permutation distribution: **SR** Knepp (block = 0.002, major triad = 0.034), **SR** Boothby (major triad = 0.022). **Faith’s PD** Knepp (major triad = 0.012, residual = 0.022), **Faith’s PD** Boothby (residual =0.014). **SES_MPD_** Knepp (block = 0.020). **SES_MNTD_** Knepp (residual = 0.012).

*2023 variance components analysis*

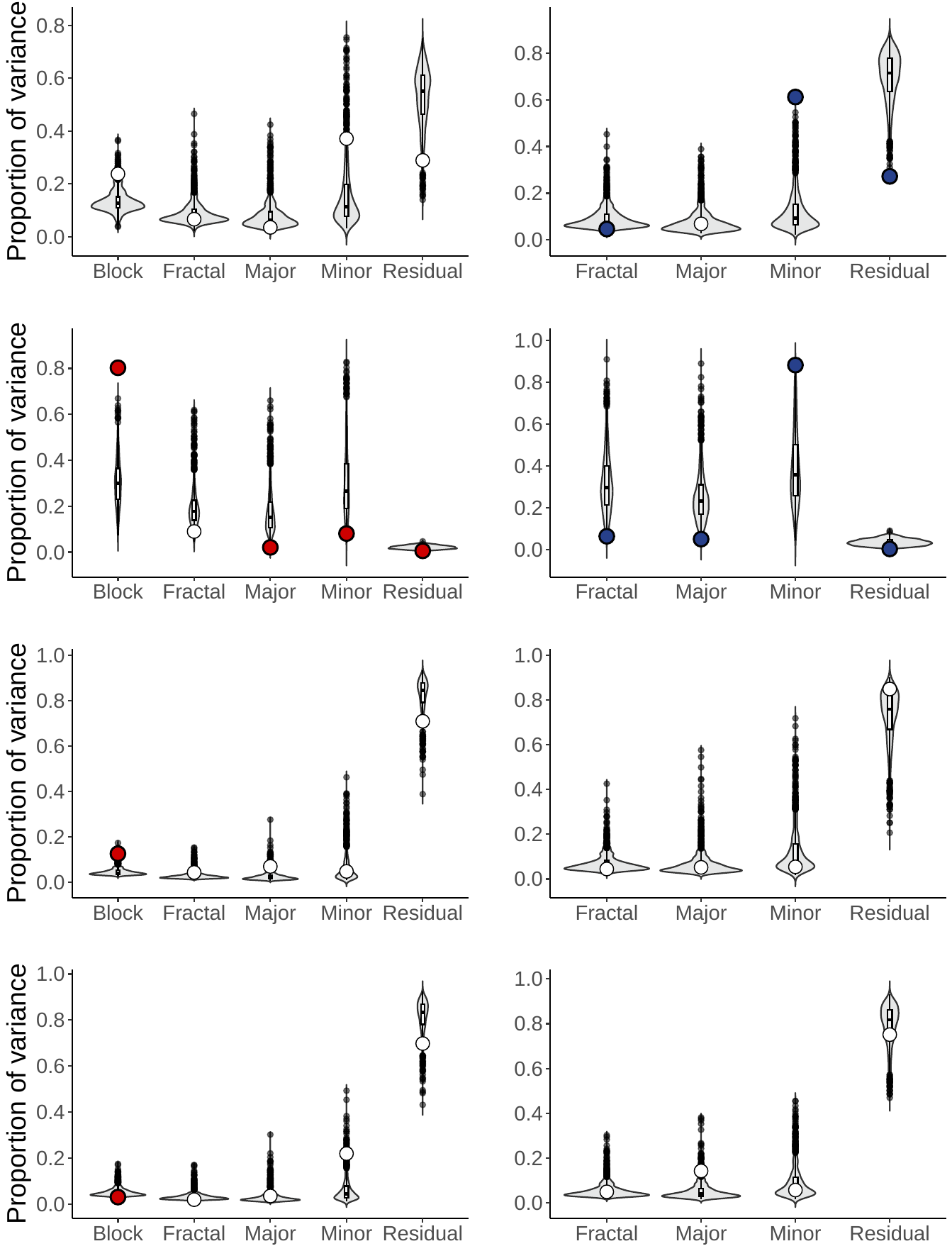

**Knepp**

**Boothby**

**Species Richness**

**Faith’s PD**

**SES_MPD_**

**SES_MNTD_**

**Figure S3.2 Pairs of variance components analysis outputs for four α-diversity metrics in 2023, showing contrasts in β-diversity between a rewilded system (the Knepp Estate) and a baseline system (Boothby Wildland).** p-values of results that are significantly different from the mean of the null permutation distribution: **SR** Boothby (fractal = 0.036, minor triad = 0.000, residual = 0.002). **Faith’s PD** Knepp (block = 0.002, major triad = 0.002, minor triad = 0.024, residual = 0.012), **Faith’s PD** Boothby (fractal = 0.006, major triad = 0.008, minor triad = 0.000, residual =0.002). **SES_MPD_** Knepp (block = 0.022). **SES_MNTD_** Knepp (residual = 0.004).

*2024 variance components analysis*

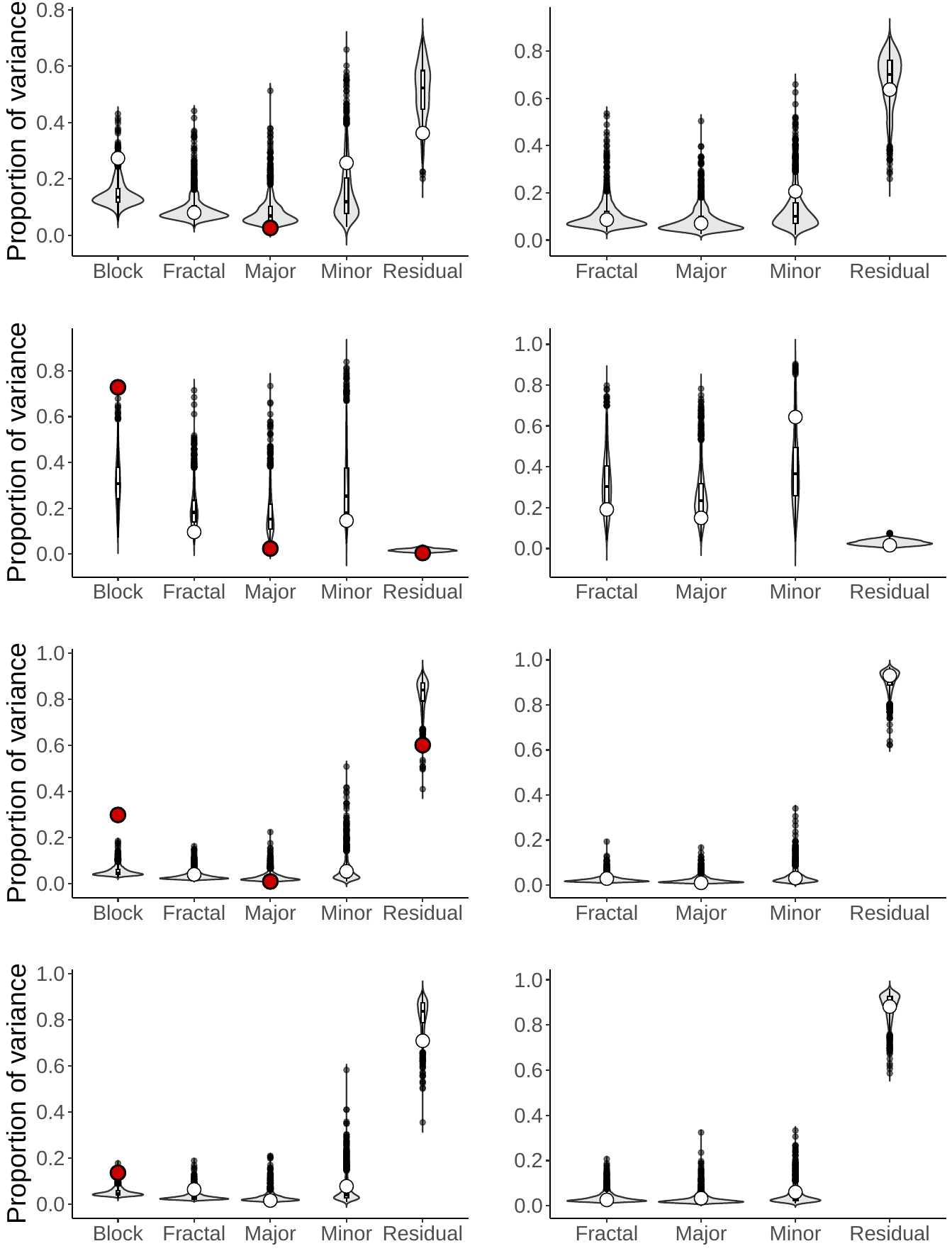

**Knepp**

**Boothby**

**Species Richness**

**Faith’s PD**

**SES_MPD_**

**SES_MNTD_**

**Figure S3.3 Pairs of variance components analysis outputs for four α-diversity metrics in 2024, showing contrasts in β-diversity between a rewilded system (the Knepp Estate) and a baseline system (Boothby Wildland).** p-values of results that are significantly different from the mean of the null permutation distribution: **SR** Knepp (major triad = 0.014). **Faith’s PD** Knepp (block = 0.002, major triad = 0.002, residual = 0.016). **SES_MPD_** Knepp (block = 0.002, major triad = 0.002, residual = 0.026). **SES_MNTD_** Knepp (block = 0.012).

**Section 4**

We expected to find a correlation between species richness and Faith’s PD. This was confirmed by correlation tests: method = Spearman

**2022** **Knepp**	rho = 0.878, S = 868, p-value <0.001

**2022** **Boothby**	rho = 0.864, S = 11144, p-value <0.001

**2023 Knepp**	rho = 0.775, S = 14608, p-value <0.001

**2023 Boothby**	rho = 0.937, S = 6720, p-value <0.001

**2024** **Knepp**	rho = 0.828, S = 11646, p-value <0.001

**2024 Boothby**	rho = 0.787, S = 16203, p-value <0.001
